# Habitat determines convergent evolution of cephalic horns in vipers

**DOI:** 10.1101/2021.10.24.465115

**Authors:** Theo Busschau, Stephane Boissinot

## Abstract

Phenotypic convergence of traits in similar environments can provide insights into the evolutionary processes shaping trait evolution. Among squamate reptiles, horn-like cephalic appendages have evolved under various selective pressures, including selection for defence, crypsis or sexual selection. Yet, among snakes, particularly vipers, the functional and evolutionary significance of horns are unknown. We used a comparative phylogenetic approach with habitat and diet data on 263 viper taxa to shed light on the selective pressures underlying horn evolution in vipers. We detected significant correlations with habitat but not diet. The relative positions of horns are ecologically divergent in that supranasal horns are positively correlated with terrestrial forest habitats while supraocular horns are negatively correlated with terrestrial forest habitats and associated with arboreal or sparsely vegetated habitats. Multiple independent origins of supranasal or supraocular horns in similar habitats provide evidence of adaptive convergence. Comparisons with other snake lineages suggest that cephalic appendages may have evolved under selection for crypsis in ambush foraging snakes.

## Introduction

Understanding the adaptive significance of phenotypic variation is an important goal in evolutionary biology. The convergence of seemingly similar traits shared among distantly related species may provide insights into the underlying evolutionary processes (Mahler *et al*., 2017). If a trait evolved independently and is associated with a similar environment or ecological niche, it is likely to be adaptive to that environment (Losos, 2011).

Among squamate reptiles, horn-like cephalic appendages evolved under various selective pressures. For example, in horned lizards, *Phrynosoma* spp., cranial horns evolved as defensive weapons against predation (Bergmann & Berk, 2012). In chameleons, horns are used in male combat, territorial displays, or to persuade females during copulation and are thus under strong sexual selection (Karsten *et al*., 2009; Stuart-fox, 2013). Rostral appendages in agamids are thought to be under sexual selection and natural selection for crypsis (Karunarathna *et al*., 2020).

In comparison to other squamates, cephalic appendages are rare among extant ophidians and their functional significance still largely unknown. The only study investigating the function of cephalic appendages in snakes is on the aquatic *Erpeton tentaculatum* Lacépède, 1800 (Homalopsidae), where nasal tentacles have a mechanosensory function to ambush fish (Catania, Leiten, & Gauthier, 2010). Among terrestrial snakes, horn-like cephalic appendages (hereafter, horns) are concentrated within the Viperidae, in which they are formed by either a single or multiple epidermal scales above the eyes (supraocular) or nasals (supranasal; Figure 1). Horns within the Viperidae are thus highly variable and occur in a variety of habitats across the globe, from sandy and rocky deserts (e.g., *Bitis caudalis* Smith, 1839) to tropical forests in terrestrial (e.g., *Bitis rhinoceros* Schlegel, 1855) and arboreal microhabitats (e.g., *Atheris ceratophora* Werner, 1896).

**Figure 1.**
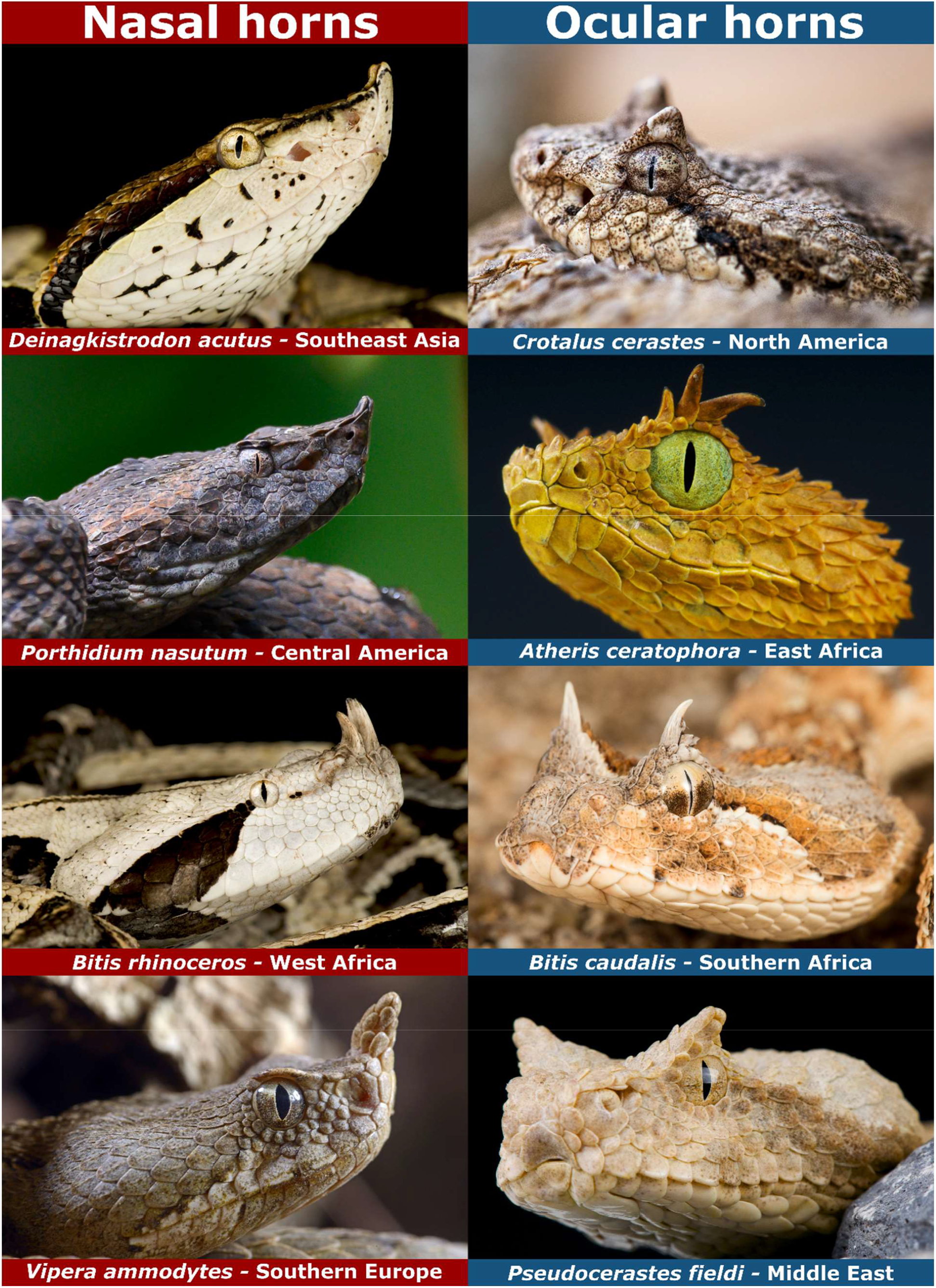
Diversity of cephalic appendages in the Viperidae. Showing examples of taxa with supranasal (nasal) and supraocular (ocular) horns along with the broad geographic region where these taxa occur. Images sourced from Adobe Stock.

Early herpetologists have speculated that supraocular horns in the horned rattlesnake, *Crotalus cerastes* Hallowell, 1854, may be of little evolutionary significance while others suggested a protective role as an eyelid when snakes enter rodent burrows (Cohen & Myres, 1970). These authors alluded to similar desert taxa inhabiting rodent burrows or rock crevices but also proposed that the enlarged supraocular scales of arboreal taxa may serve a protective function among dense foliage. Wagner & Wilms (2010) hypothesized that cephalic horns among vipers may serve a role in sexual recognition as in other squamates.

More recently, several hypotheses have been proposed in non-peer reviewed literature. Naish (2009) suggested horns may aid in camouflage of forest-living species by obscuring snakes’ silhouette, but due to the diversity of cephalic appendages and wide range of ecologies horns may have evolved for multiple reasons. del Mármol (2020) supported the camouflage hypothesis in desert taxa and further speculated that horns may act as lures to attract rodent prey in sandy deserts or serve a protective role during prey capture.

Despite wide speculation, the evolution of cephalic horns in Viperidae has not gained any attention in recent scientific literature. Here, we assembled habitat and diet information on 263 viper taxa and use a phylogenetic comparative approach to shed light on the selection pressures underlying horn evolution. By assessing correlation between horns and habitat or diet, we tested the hypothesis that the presence of horns exhibits clear ecological association and evolved repeatedly in similar environments.

## Methods

### Data collection

As the basis of our comparative analyses, we used a previously published time-calibrated phylogeny derived from six mitochondrial and five nuclear genes for 263 Viperidae taxa (Alencar *et al*., 2016). For each taxon in the phylogeny, we assembled ecological and morphological trait data from published literature and online databases. The data was coded in a binary matrix, recording the presence/absence of a taxon under each variable. Habitat categories were defined by the IUCN habitat classification scheme (Version 3.1; https://www.iucnredlist.org/resources/habitat-classification-scheme), comprising forest, savanna, shrubland, grassland and desert habitats. We further recorded the presence in five microhabitat categories, including sandy (e.g., windblown sand dunes), rocky (e.g., outcrops, karst, cliffs), arboreal (including semi-arboreal), terrestrial forest (fully terrestrial taxa in forest habitat), and water-associated (e.g., swamps, marshes, rivers). We were able to obtain diet information for 224 out of the 263 taxa where we recorded the presence of six prey classes (mammals, reptiles, birds, amphibians, fish, and invertebrates) known from the diet of each taxon. Morphological traits were recorded from images on iNaturalist (https://www.inaturalist.org/), The Reptile Database (Uetz *et al*., 2021), or species descriptions for taxa where images were not available. We recorded the presence of ocular or nasal horns, defined as tapered projections above the eyes or rostral, respectively (Figure 1). We also identified intermediate characters that comprise either a pronounced ridge or comparatively smaller projections that do not taper to a point (Figure S1). We did not consider dorsally flattened ocular ridges (e.g., large supraoculars of *Crotalus* spp. or *Vipera* spp.) to be intermediate character states of ocular horns. We were unable to account for intraspecific polymorphism of character states due to the binary nature of the correlation analyses employed in this study, but instead prioritized the maximal character states that may be present in a taxon. All binary statistical analyses were performed on two datasets. The ‘horns’ dataset comprises only the presence/absence of horns, and the ‘characters’ dataset comprises the presence/absence of horns or intermediate characters. In addition to analyzing the combined ‘horns’ and ‘characters’ datasets, we also analyzed nasal and ocular traits separately. All statistical analyses were conducted in R v.4.0.2 (R Core Team, 2020). The data underlying this work will be made available with the published version of the manuscript.

### Ancestral state reconstruction

To determine the most parsimonious models of evolution, we fit various continuous-time Markov models to our datasets, using the ‘fitMK’ function in the R package *phytools* v.0.7.80 (Revell, 2012). For multiple states, e.g., ocular horns, nasal horns, or no horns, we fit equal-rates (ER), symmetric-rates (SYM), and all-rates-different (ARD) models. Additionally, for the multistate datasets including horns and intermediate characters, we fit the ER, ARD and SYM models along with ordered versions of each model where a trait needs to transition to an intermediate state before transitioning to the horned state. The best-fit models were selected using Akaike Information Criterion (AIC) and AIC weights (AICw). Ancestral state reconstruction of horn evolution was then estimated with the best-fit models (Table S1) using stochastic character mapping, simulating 1000 maps using the ‘make.simmap’ function in *phytools*. The root states (pi) of all models were estimated from the stationary distribution of the transition matrix (Q) assuming a flat prior.

We tested for phylogenetic signal using *D* statistics (Fritz & Purvis, 2010) in the R package *caper* v.1.0.1 (Orme *et al*., 2013). The *D* statistic is a measure of phylogenetic clumping where *D* is equal to 1 if a binary trait is randomly distributed across the phylogeny, 0 if the trait is phylogenetically clumped as if it had evolved by Brownian motion, and exceeds this range if a trait is overly dispersed or more clumped than expected by Brownian motion (Fritz & Purvis, 2010).

### Habitat association

To test for correlations between association with habitat and the presence of ‘horns’ or ‘characters’, we employed two complementary approaches in *phytools*. First, we used the ‘fitPagel’ function that fits a model of correlated evolution between two binary traits (Pagel, 1994). This method uses a likelihood ratio test to compare two models in which binary traits either evolve independently or in a dependent manner. In our analyses trait states were set as the dependent variable and the “fitDiscrete” method was used to fit the model of discrete character evolution. Since vipers with ocular horns occur in terrestrial and arboreal habitats, we performed the analyses twice on each dataset: First on all taxa and then by removing the 75 arboreal taxa to identify terrestrial habitats where the selection pressures underlying horn evolution may be similar to arboreal microhabitats. The threshold for declaring statistical significance was set to p < 0.05, and p-values were corrected for multiple comparisons using False Discovery Rate (FDR) correction.

For significant dependent models of evolution, we quantified the correlation between habitat and traits using a Bayesian version of the quantitative threshold model (Felsenstein, Ackerly, & Mcpeek, 2012) in the function ‘threshBayes’. We ran four independent chains for 10 million generations, sampling every 2000, and discarding the first 20% of each chain as burn-in. We summarized the 90% highest posterior density (HPD) to assess significance in the probability of direction (PD), an index of effect existence (Makowski *et al*., 2019). The effect is considered significant if the entire 90% HPD is either positive or negative, i.e., the PD > 95%.

### Relative diet specificity

First, we tested for correlated evolution between prey class and ‘horns’ or ‘characters’ using ‘fitPagel’ as above, which did not yield significant correlations (Table S2). Therefore, we explored whether traits may be associated with specificity to a particular prey class using a diet specificity index comparable to that of Maritz *et al*. (2016). We transformed the presence/absence data to a relative diet specificity index (DSi) for each of the six prey classes, which is presence (1) or absence (0) divided by the number of prey classes in the diet of a taxon and ranked from zero. Thus, zero represents no consumption and higher rank represents higher relative specificity to a particular prey class. Although this scaled index does not provide an estimation of prey specificity within taxa, it can with the available data provide a relative comparison among taxa. Only six taxa were recorded eating fish, therefore this prey class was not included in further analyses. We tested for a correlation between traits and diet specificity using phylogenetic generalized least squares (PGLS), implemented with residual randomization in a permutation procedure in the *RRPP* package (Collyer & Adams, 2018) with 10000 permutations for significance testing. The multivariate response variables (DSi for five prey classes) were evaluated using MANOVA and Pillai’s trace test statistic, followed by univariate ANOVAs to assess differences in specificity to each prey class. To test for dietary differences between trait states associated with different habitats, we included an interaction term between trait state and habitat. We specifically tested interaction among correlated trait and habitat variables.

## Results

We identified 35 taxa out of 263 in which horns are present (13.3%), 14 (5.3%) with nasal horns and 21 (8%) with ocular horns (Figure 2). Additionally, we identified 33 taxa in which intermediate character states were present (12.5%), 22 (8.4%) nasal and 11 (4.2%) ocular. In total we recorded 59 taxa (22.4%) in which characters were present, 36 (13.7%) with nasal characters and 32 (12.2%) with ocular characters. Of these, 9 taxa (3.4%) can have both nasal and ocular characters. The binary analyses were performed separately on the ‘horns’ dataset, including only the presence/absence of horns (35 taxa), and on the ‘characters’ dataset, including the presence of horns and intermediate characters (59 taxa).

**Figure 2.**
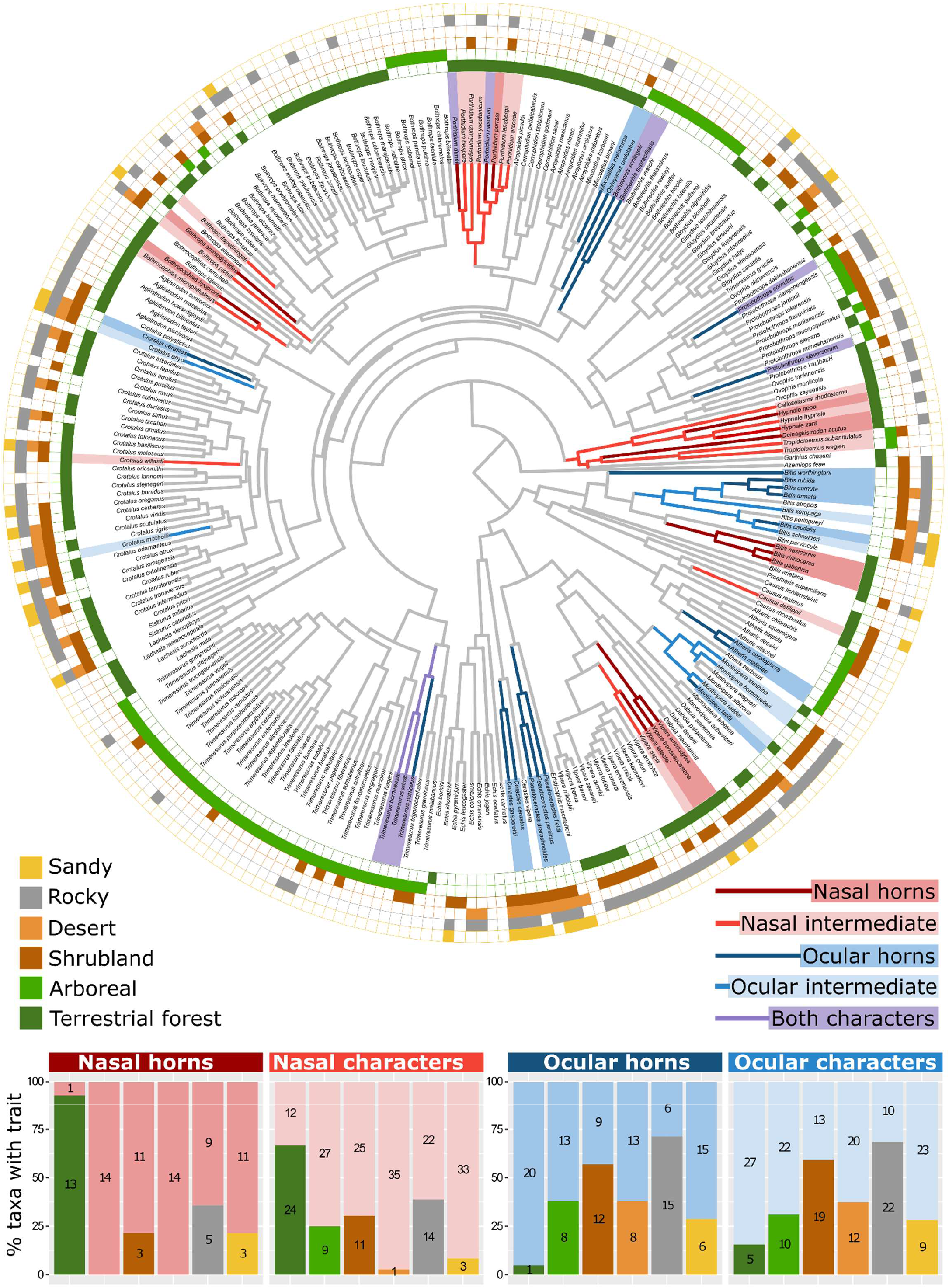
Evolutionary history of cephalic horns in the Viperidae along with correlated habitat variables. Branches are colored to show the character states estimated with the highest probability at each node. Exact trait probabilities are available in Figures S2-S5. Branch colors prioritize the full horn character states if present. Shading of taxon labels depict the character states of each taxon and the binary matrix indicates the association with each habitat variable. Bar graphs show the relative proportion of taxa with each trait associated with the correlated habitat variables compared to the proportion of taxa with the trait not associated with each habitat. Values in the columns depict the absolute number of taxa. Note that ‘characters’ include horns and intermediate character states.

### Ancestral state reconstruction

The posterior distribution of 1000 stochastic character maps on the ‘horns’ dataset (Figure S2) estimated an average of 14 independent origins of ocular horns (95% HPD: 12-16) with 1 reversal (95% HPD: 0-3) and 11 origins of nasal horns (95% HPD: 9-13) with 1 reversal (95% HPD: 0-3). The posterior distribution on the ‘characters’ dataset (Figure S3) estimated 12 origins of ocular characters (95% HPD: 10-15) and 4 reversals (95% HPD: 1-7); 12 independent origins of nasal characters (95% HPD: 9-14) with 2 reversals (95% HPD: 0-4) and 3 transitions to a state with both characters (95% HPD: 2-4). There was a mean of 4 independent origins of a state with both characters from a state with no characters (95% HPD: 3-5).

For the multistate ocular and nasal datasets, the ordered models of evolution, where a trait needs to transition to an intermediate state before transitioning to the horned state, generally retrieved lower AIC values compared to the respective unordered models (Table S1). For both nasal and ocular horns, the ancestral state reconstruction implementing the best-fit ordered models estimated multiple independent origins of intermediate characters followed by frequent transitions between intermediate characters and horns, and less frequent reversals back to a state with no characters (Figure 3; Figures S4 & S5). *D* statistics for all datasets were significantly different from 1 and not significantly different from 0 (Table S3), suggesting traits are evolving as expected under Brownian motion. The estimated values of the *D* statistic generally suggest stronger phylogenetic signal in the ‘characters’ datasets (characters: *D* = -0.06; ocular: *D* = 0.08; nasal: *D* = -0.25) compared to the ‘horns’ datasets (horns: *D* = 0.33; ocular: *D* = 0.16; nasal: *D* = 0.18).

**Figure 3.**
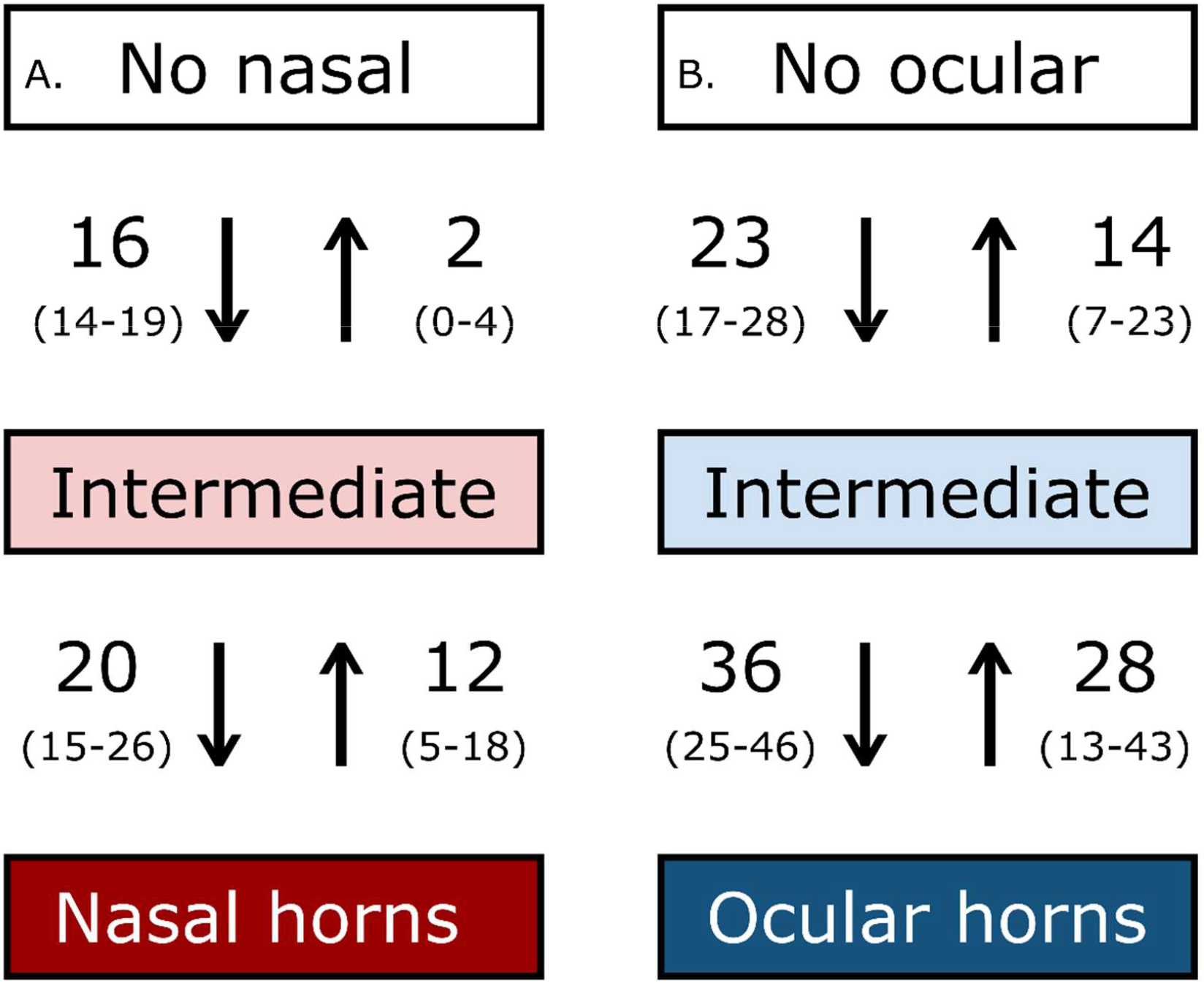
Mean number of transitions between nasal (A) and ocular (B) character states estimated from the posterior distribution of 1000 stochastic character maps (Figures S4 & S5). 95% highest posterior density interval shown in parentheses.

### Habitat association

Habitat dependent models of evolution for the combined ‘horns’ or ‘characters’ datasets were not significantly better compared to the independent models where traits and habitat association evolve independently (Table S4). However, for both ocular and nasal horns the dependent models performed better where the rate of evolution was dependent on a taxon’s association with terrestrial forest habitats (Tables 1 & S4A). When arboreal taxa were removed to detect correlations among terrestrial habitats, the models where ocular horns depend on terrestrial forest, shrubland, rocky and desert habitats performed better compared to the respective independent models (Tables 1 & S4A). Correlating the presence of horns with a taxon’s occurrence in these habitats under the threshold model retrieved a moderate but significantly positive correlation between nasal horns and terrestrial forest habitats (r= 0.38; PD > 95%) while ocular horns had a moderate but significant negative correlation with terrestrial forest habitats (r= -0.45; PD > 95%; Figures 4 & S6). After removing arboreal taxa, the correlations between ocular horns and occurrence in shrubland (r= 0.35; PD > 95%), rocky (r= 0.35; PD > 95%) and desert (r= 0.33; PD > 95%) habitats were moderate but significantly positive (Figure S6).

**Table 1.** log-likelihood (logL) for models of evolution between habitat and horns within the Viperidae. Showing p-values and FDR corrected p-values for independent models of evolution based on the likelihood ratio test. Significant p-values shown in bold. Complete results on all datasets are available in Table S4.

|  | Nasal horns |  |  |  | Ocular horns excluding arboreal taxa |  |  |  |
| --- | --- | --- | --- | --- | --- | --- | --- | --- |
| Habitat | Independent | Dependent | p-value | Corrected | Independent | Dependent | p-value | Corrected |
|  | logL | logL |  | p-value | logL | logL |  | p-value |
| forest | -164.508 | -163.599 | 0.403 | 0.832 | - | - | - | - |
| **terrestrial forest** | -195.466 | -190.466 | **0.007** | 0.067* | -144.187 | -136.615 | **0.001** | **0.004** |
| arboreal | -112.551 | -112.551 | 1.000 | 1.000 | - | - | - | - |
| savanna | -148.534 | -148.061 | 0.623 | 0.832 | -124.588 | -124.194 | 0.675 | 0.771 |
| **shrubland** | -198.307 | -197.329 | 0.376 | 0.832 | -151.070 | -145.660 | **0.004** | **0.012** |
| grassland | -188.919 | -188.512 | 0.666 | 0.832 | -153.892 | -152.360 | 0.216 | 0.288 |
| sandy | -125.469 | -125.011 | 0.633 | 0.832 | -110.996 | -108.243 | 0.064 | 0.102 |
| **rocky** | -188.286 | -187.357 | 0.395 | 0.832 | -146.751 | -140.937 | **0.003** | **0.012** |
| **desert** | -125.146 | -125.146 | 1.000 | 1.000 | -114.829 | -110.771 | **0.017** | **0.035** |
| water-associated | -125.146 | -200.215 | 0.490 | 0.832 | -138.649 | -138.649 | 1.000 | 1.000 |

\*Corrected p-value not significant at alpha = 0.05 but correlation under the threshold model is significant (Figure 4).

**Figure 4.**
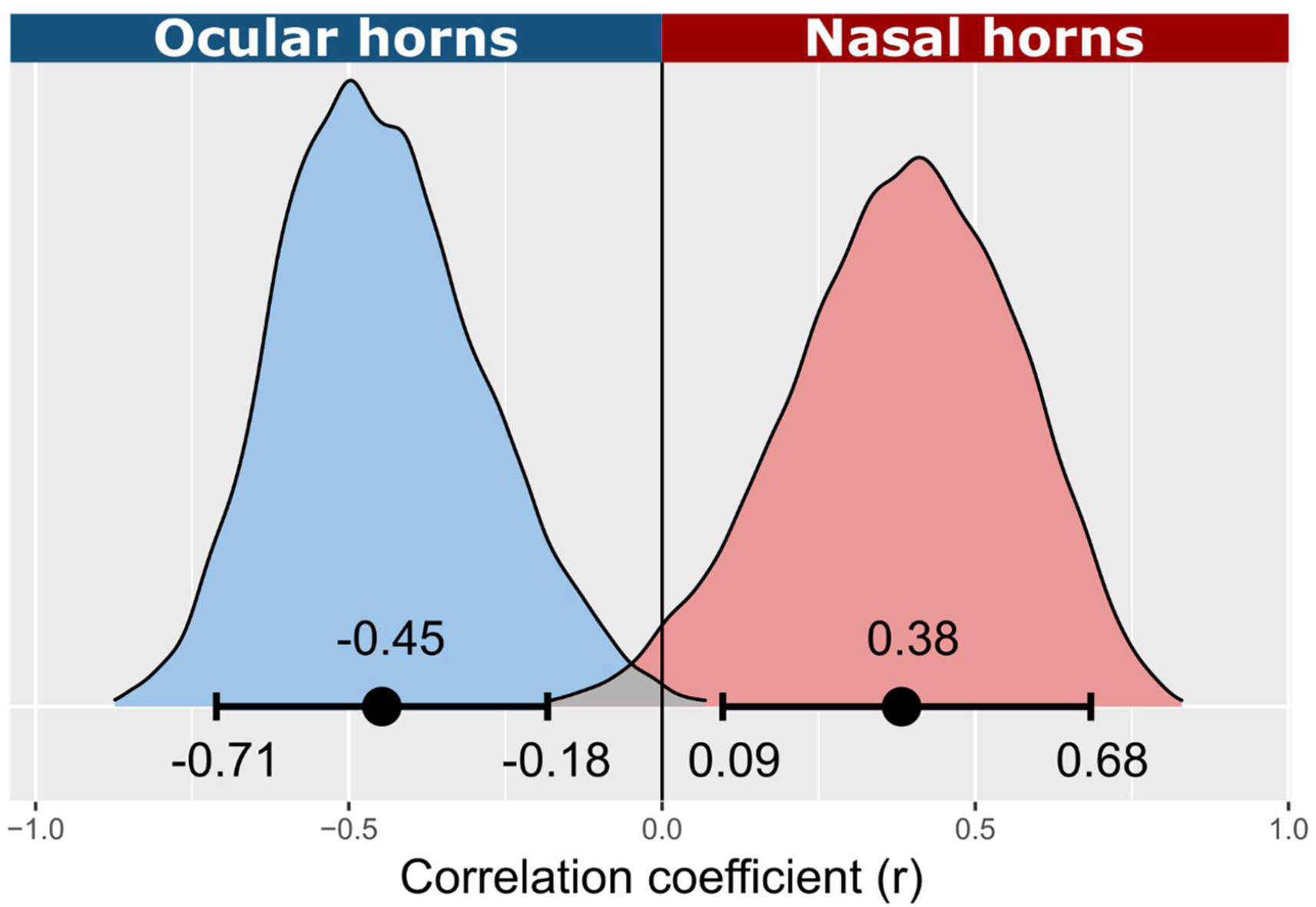
Contrasting correlation between horns and terrestrial forest habitat. Showing posterior estimate of the correlation coefficient (r) between the presence of nasal or ocular horns and terrestrial forest habitat under the threshold model. Dot indicates the mean of the posterior distribution. Error bars indicate the 90% highest posterior density, i.e., the probability of direction is >95%.

As with the ‘horns’ dataset, the correlation between ocular characters and terrestrial forest was moderate but significantly negative (r= -0.39; PD > 95%; Figure S7). In contrast, for nasal characters, the dependent model of evolution was not significantly better than the independent model (Table S4B) and assessing the correlation between forest habitats and nasal characters under the threshold model revealed no significant correlation (Figure S7). When arboreal taxa were removed, the rate of ocular character evolution was dependent on the association with terrestrial forest, shrubland, rocky, and desert habitats, as well as sandy habitat (Table S4B). Under the threshold model, the correlation between ocular characters and terrestrial forest (r = -0.41; PD > 95%), shrubland (r= 0.31; PD > 95%), rocky (r= 0.38; PD > 95%) and desert habitats (r= 0.40; PD > 95%) were significant but the correlation with sandy habitat was not significantly positive (r = 0.30; 90% HPD: -0.02 – 0.63; Figure S7).

Taxa with nasal horns exhibit the highest shared association with terrestrial forest microhabitats, 13 out of 14 taxa (92.9%; Figure 2). Similarly, despite not detecting significant correlations, the highest shared habitat association among taxa with nasal characters is with forest habitat (33/36; 91.6%), followed by terrestrial forest microhabitats (24/36; 66.6%).

Only one out of 21 taxa with ocular horns is associated with terrestrial forest microhabitats (4.5%), reflecting shared dissociation. Among the habitats correlated with ocular horns, the highest shared association is with rocky microhabitats (15/21; 71.4%), followed by shrubland (12/21; 57,1%), desert (8/21; 38.1%), arboreal (8/21; 38.1%) and sandy (6/21; 28.6%). This trend is consistent for taxa with ocular characters (Figure 2).

### Relative diet specificity

The presence of horns or characters did not have a significant effect on relative diet specificity, nor did the different horn or character types (Tables S5A & S6A). The interaction terms between traits and habitat generally did also not have significant effects on relative diet specificity, yet across all models the habitat variables had significant multivariate or univariate effects (Tables S5 & S6). The interaction term between arboreal microhabitats and ocular traits did have a significant effect on relative specificity to invertebrate prey. Arboreal taxa with ocular traits had significantly lower relative specificity compared to arboreal taxa without ocular traits (Tables S5F & S6E). There are, however, only six out of 75 arboreal taxa recorded feeding on invertebrates (although none with ocular traits), thus this result should be interpreted with caution. The interaction term between ocular horns and desert habitat had a significant multivariate effect on relative diet specificity, where diet specificity of taxa with ocular horns associated with deserts is significantly different from taxa with ocular horns not associated with deserts (Table S5G). This interaction term also had a significant univariate effect on relative specificity to reptile prey, with no significant pairwise differences, as well as a significant effect on relative specificity to bird prey, where taxa with ocular horns in deserts had significantly higher specificity compared to all other groups (Table S5G). However, in the ‘characters’ dataset the interaction term between desert and ocular characters had a significant effect only on relative specificity to bird prey, where taxa with ocular characters in deserts have significantly higher specificity compared to taxa with ocular characters not associated with deserts (Table S6F).

There is a notable difference in the proportion of taxa with each trait that have been recorded feeding on amphibians (Figure S8). 78.6% and 69.4% of taxa with nasal horns and characters were recorded feeding on amphibians compared to only 28.6% and 31.3% of taxa with ocular characters, while the proportion of taxa feeding on other prey classes were comparable. This reflects the general occurrence of taxa with nasal horns in mesic forest environments compared to taxa with ocular horns in arid environments.

## Discussion

Our analyses revealed significant correlations between the evolution of horns and habitat despite the low frequency of Viperidae taxa in which horns and intermediate characters are present. The relative positions of horns are ecologically divergent in that taxa with nasal characters are associated with terrestrial forest and forest habitats while taxa with ocular characters are dissociated with terrestrial forest habitats and associated with arboreal or sparsely vegetated terrestrial habitats (Figures 2 & 4). Furthermore, multiple independent origins of similar traits in similar habitats across the globe provides evidence of adaptive convergence.

Multiple origins and transitions between character states suggest that cephalic horns are highly labile within the Viperidae. Frequent transitions, particularly between intermediate character states and horns (Figure 3), fit a theoretical model of evolution under Brownian motion whereby a continuously varied trait fluctuates randomly over time (Revell, Harmon, & Collar, 2008). This observation is supported by significant Brownian phylogenetic structure across all datasets (Table S3). Evolution by Brownian motion does not necessarily imply random drift. Instead strong phylogenetic signal and oscillation in the extent of character states can also be explained by fluctuating selection in variable environments over time or among populations (O’Meara *et al*., 2006; Revell *et al*., 2008; Bell, 2010). In this regard, reduced rate of fluctuating selection, stabilizing or divergent selection can decrease the strength of phylogenetic signal (Revell *et al*., 2008). Therefore, the ordered models of evolution along with a reduction in phylogenetic signal between the ‘characters’ and ‘horns’ datasets may suggest that horns evolved under selection from an intermediate state. Furthermore, the non-random phylogenetic clumping of ocular and nasal characters in particular habitat types suggest that characters are selected for in the habitats with which they associate or selected against in the habitats with which they do not associate while evolving randomly in other habitats, or both (Figure 2). Correlations with habitat make it tempting to assume that the observed patterns of horn evolution was driven by frequent habitat shifts throughout the planet’s dynamic history. It should, however, be noted that vipers, including polymorphic horned taxa, may exhibit substantial phenotypic variability among and within populations (Wagner & Wilms, 2010; Pizzigalli *et al*., 2020). It is therefore likely that microevolutionary processes at the population level contribute to the frequent transitions of character states observed at a macroevolutionary scale (Li *et al*., 2018).

Significant correlations between horns and habitat may not be sufficient to determine causation of horn evolution in vipers, especially since < 13.3% of the vipers in our analyses possess horns, although habitat use has been attributed to distant morphological convergence in other snake families (Grundler & Rabosky, 2014; Esquerré & Scott Keogh, 2016). Furthermore, our inability to detect correlations between diet and horns is to be expected. As ambush predators vipers are generally opportunistic with wide diet breadths (Glaudas *et al*., 2019), thus diet is likely to be constrained mainly by available prey in the environment without limiting morphological convergence (Grundler & Rabosky, 2014). This notion is evident in the PGLS where habitat had the largest effect on relative diet specificity (Tables S5 & S6), and the significant differences detected between ocular characters associate with deserts and ocular characters not associated with desert habitat (Tables S5G & S6F).

Ambush predators are known for using cryptic phenotypes to avoid detection, ensuring that prey move within striking distance to improve hunting success (Pembury Smith & Ruxton, 2020). Like vipers, the few snake lineages with cephalic appendages also employ ambush foraging. Nasal appendages occur in *Ahaetulla* (Colubridae) and *Langaha* (Lamprophiidae) species, two distantly related arboreal genera thought to rely on crypsis to ambush diurnal prey (Henderson & Binder, 1980; Kuchling, 2003; Tingle, 2012; Mallik *et al*., 2020). Two of the three *Langaha* species possess both nasal and ocular appendages (Kuchling, 2003), which is comparable to the arboreal viper taxa we recorded with both characters (Figure 2). Nasal appendages also occur in two arboreal forest-living *Gonyosoma* (Colubridae) species (Peng *et al*., 2021). Information on the foraging behavior of this genus is limited but they may use ambush and active foraging modes (Hodges, 2019). In non-arboreal habitats, *Trachyboa boulengeri* Peracca, 1910 (Tropidophiidae) possesses both characters, is highly cryptic and utilizes an ambush technique to prey on fish and frogs (Dieter Lehmann, 1969; Arteaga, 2021). Ambush foraging has also been observed in the horned sea snake, *Hydrophis peronii* Duméril, 1853 (Elapidae; Borsa, 2008). *Acanthophis* (Elapidae), in which some species have horn-like supraocular scales, share remarkable morphological convergence with terrestrial vipers and like many vipers also employ caudal luring to attract prey (Shine, Spencer, & Keogh, 2014; Crowe-Riddell *et al*., 2021). These comparisons may support the hypothesis that horns evolved under selection for crypsis in ambush foraging snakes.

Crypsis is highly dependent on habitat and hunting location of predators, which often results in different modes of camouflage among related organisms in forested compared to open habitats (Pembury Smith & Ruxton, 2020; Caro & Koneru, 2021). This trend is apparent in the Viperidae, especially among *Bitis* species where terrestrial forest taxa possess nasal horns and open habitat taxa possess ocular horns (Figures 1, 2 & 4). In most cases cryptic or disruptive colors are sufficient forms of camouflage (Egan *et al*., 2016; Hughes, Liggins, & Stevens, 2019). Yet, there may be scenarios where disruption or deception by secondary appendages could reduce detection and increase fitness. For example, over evolutionary time predator or prey could adapt to the camouflage strategies of the other and thereby drive the evolution of improved crypsis (Pembury Smith & Ruxton, 2020). Crypsis can also evolve to enhance diurnal concealment or to the differential visual systems of specific organisms (Hughes *et al*., 2019; Caro & Koneru, 2021).

Comparisons between vipers, other snakes and other forest-living squamates (Losos *et al*., 2012; Breuil *et al*., 2019; Karunarathna *et al*., 2020) suggests a broader convergence of nasal appendages in forest-associated taxa. Terrestrial forest habitats are typically covered in a layer of leaf litter or foliage which terrestrial vipers utilize as concealment to ambush prey. Here, nasal horns could potentially disrupt the outline of a viper’s head, making it more difficult to detect against a background of leaves and twigs. In contrast, vipers in arboreal, rocky, or sparsely vegetated open habitats may be more limited to exposed ambush sites with little cover. It has previously been hypothesized that the vertical pupils of ambush foraging snakes aid in concealment by disrupting the round outline of the eye (Brischoux, Pizzatto, & Shine, 2010). The position of supraocular horns would contribute significantly to concealing the eyes and disrupting a viper’s silhouette in more exposed ambush sites. In this regard, the selection pressures imposed by the arboreal ambush sites of *Bothriechis schlegelii* Berthold, 1846 may be analogous to the arid rocky habitats of *Pseudocerastes* species. Both these taxa are able to rely on crypsis to ambush diurnal prey while avoiding predation (Sorrell, 2009; del Mármol, Mozaffari, & Gállego, 2016). Similarly, two ocular horned vipers occurring at high altitudes, *Mixcoatlus melanurus* Müller, 1924 and *Ophryacus undulatus* Jan, 1859, are thought to be constrained to diurnal foraging habits due to low nocturnal temperatures (O’Shea, 2018). The occurrence of supraocular horns in cryptic lizard genera, e.g., *Acanthosaura* and *Lyriocephalus* (Agamidae) as well as *Uroplatus* (Gekkonidae), may further support their role in crypsis (Rodda, 2020).

While our analyses provide evidence to support convergent evolution of Viperidae horns in similar habitats, more detailed studies are needed to fully understand their role in crypsis. The relationship between horns and camouflage may be complex, especially among variable populations and/or environments where there are likely to be community dependent tradeoffs between hunting success and predator avoidance.

## Acknowledgments

We thank Adriaan Jordaan, Courtney Hundermark, Tyrone Ping and Sterrin Smalbrugge for their assistance with the literature review, and Bryan Maritz for sharing diet data. We are grateful to Sandra Goutte for suggestions on the analyses, and Sebastian Kirchhof and Justin Wilcox for comments on the manuscript. This work was supported in part by the NYU Abu Dhabi Global PhD Student Fellowship.

## Supporting information

**Table S1**. Best-fit continuous-time Markov models.

**Table S2. ‘**fitPagel’ results on diet classes.

**Table S3**. Phylogenetic signal, *D* statistics.

**Table S4. ‘**fitPagel’ results on habitat.

**Table S5**. Phylogenetic generalized least squares on relative diet specificity. Horns dataset.

**Table S6**. Phylogenetic generalized least squares on relative diet specificity. Characters dataset.

**Figure S1**. Examples of intermediate cephalic characters in the Viperidae.

**Figure S2**. Ancestral state reconstruction of cephalic horns in Viperidae.

**Figure S3**. Ancestral state reconstruction of cephalic characters in Viperidae.

**Figure S4**. Ancestral state reconstruction of nasal horns in Viperidae.

**Figure S5**. Ancestral state reconstruction of ocular horns in Viperidae.

**Figure S6**. Posterior estimate of the correlation coefficient (r) between the presence of horns and habitat.

**Figure S7**. Posterior estimate of the correlation coefficient (r) between the presence of characters and habitat.

**Figure S8**. Relative proportion of Viperidae taxa with cephalic traits recorded feeding on each prey class.

## Notes

### Competing Interest Statement

The authors have declared no competing interest.

