## Supplementary material for "Habitat determines convergent evolution of cephalic horns in vipers": Figure S

Evolution of horns in vipers

Theo Busschau^1^ & Stephane Boissinot^1^

New York University Abu Dhabi, Saadiyat Island, Abu Dhabi, United Arab Emirates

Corresponding authors:

 (T.B.)

 (S.B.)

**Supplementary figures**

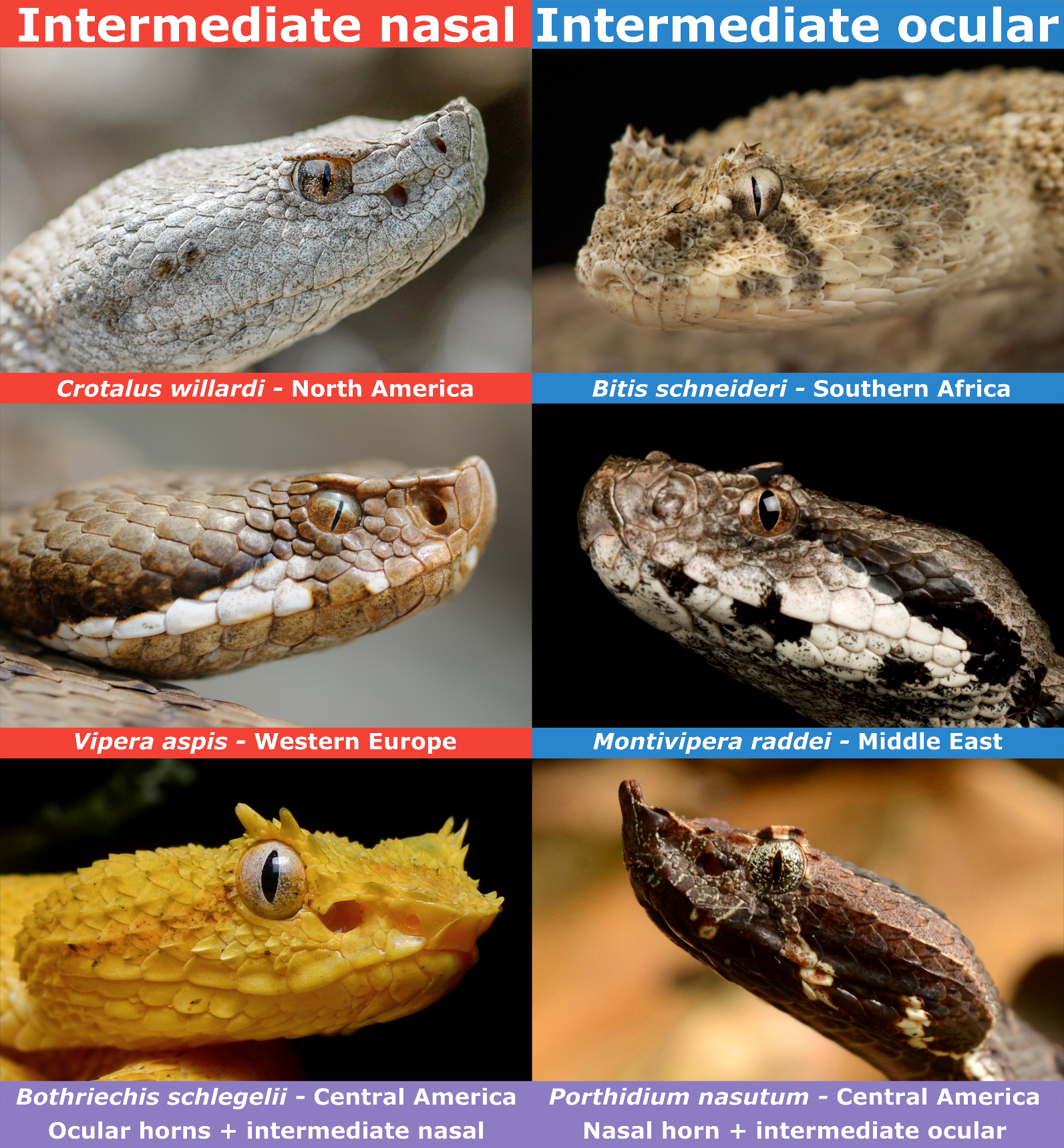

**Figure S1.** Intermediate cephalic characters in the Viperidae. Showing examples of taxa with intermediate supranasal (nasal) and intermediate supraocular (ocular) characters along with the broad geographic region where these taxa occur. Bottom two images show examples of taxa with both characters. Top four images sourced from Shutterstock. *Bothriechis schlegelii* by Geoff Gallice, used under license CC BY 2.0 (<https://creativecommons.org/licenses/by/2.0>) via Wikimedia Commons (<https://commons.wikimedia.org/wiki/File:Bothriechis_schlegelii_(La_Selva_Biological_Station).jpg>). *Porthidium nasutum* by Josue Ramos Galdamez, used under license CC BY-NC 4.0 (<https://creativecommons.org/licenses/by-nc/4.0>) via iNaturalist (<https://www.inaturalist.org/photos/121231602>). Images cropped and oriented for presentation.

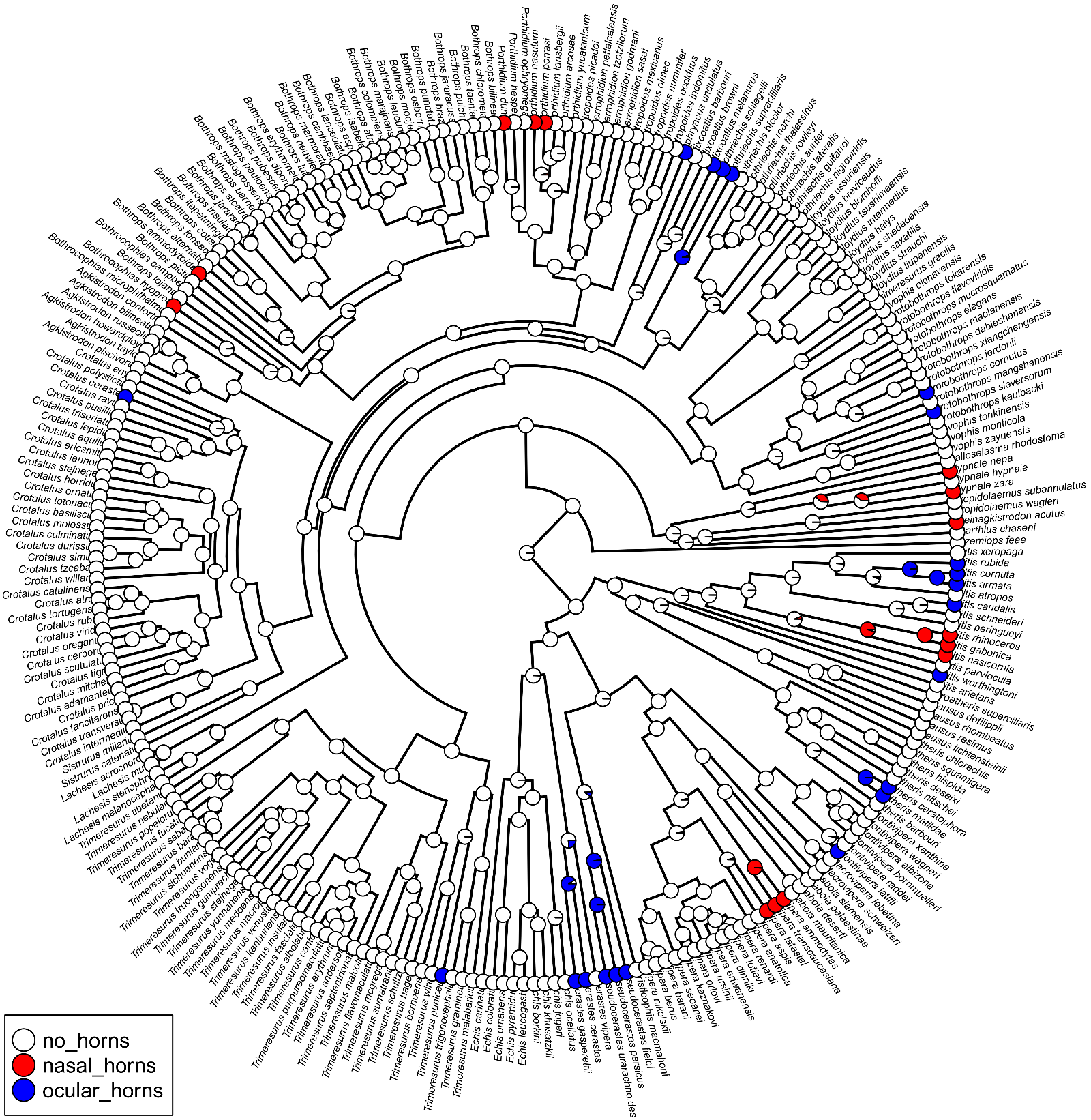

**Figure S2.** Ancestral state reconstruction of cephalic horns within the Viperidae. Pies depict the posterior probability of character states at each node obtained from 1000 stochastic character maps.

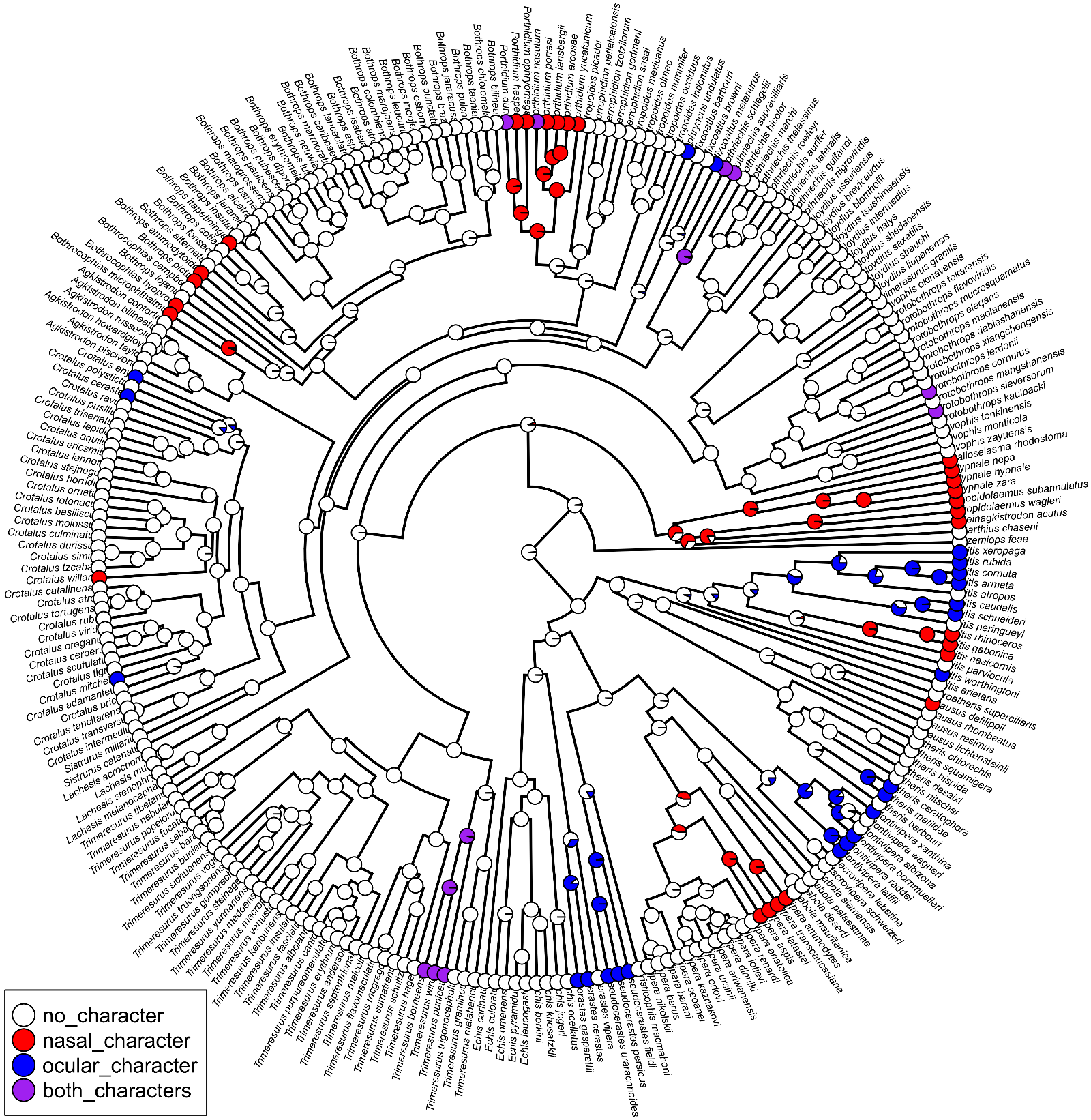

**Figure S3.** Ancestral state reconstruction of cephalic characters within the Viperidae, including horns and intermediate states as characters. Pies depict the posterior probability of character states at each node obtained from 1000 stochastic character maps.

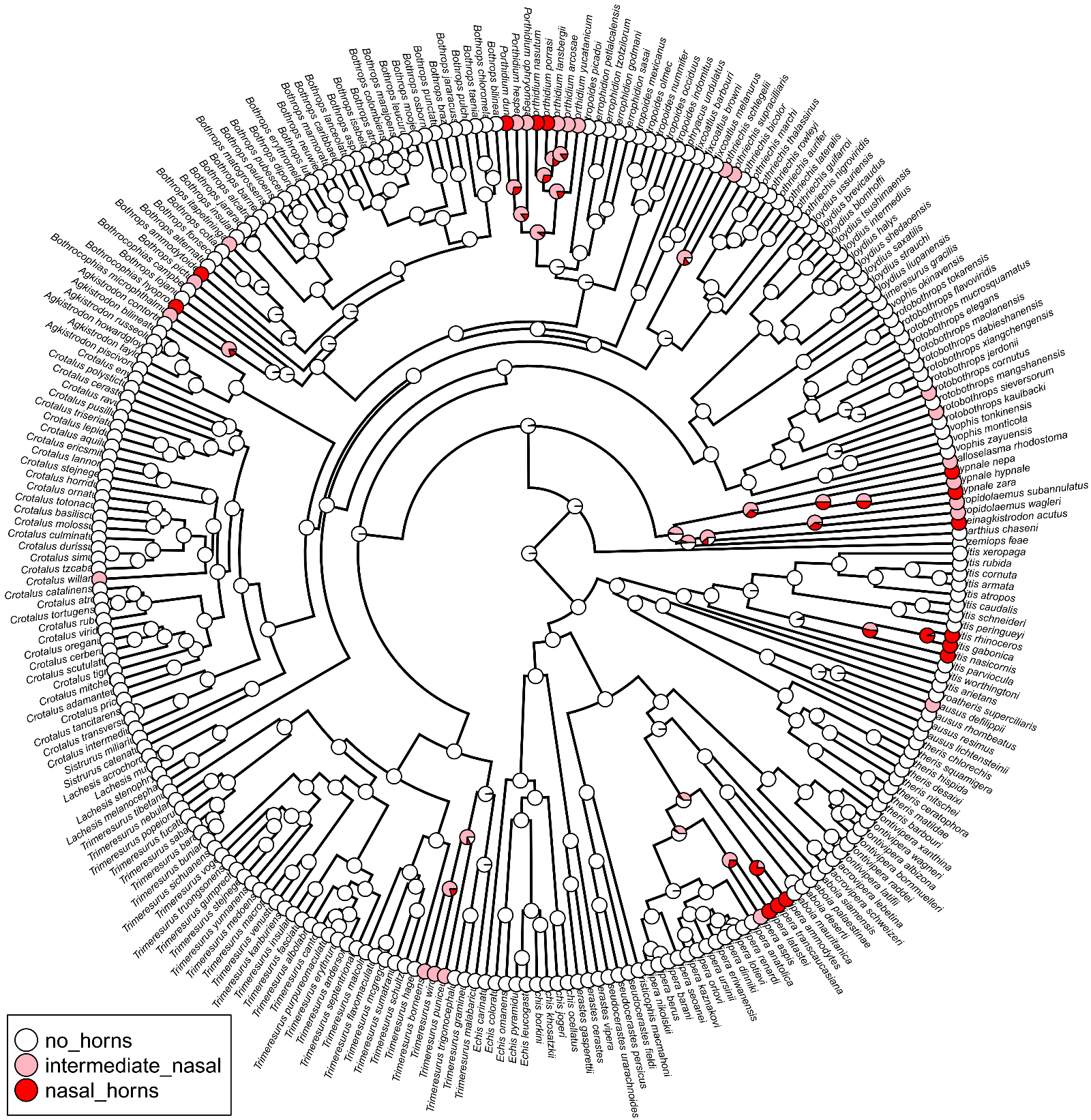

**Figure S4.** Ancestral state reconstruction of nasal horns within the Viperidae. Pies depict the posterior probability of character states at each node obtained from 1000 stochastic character maps.

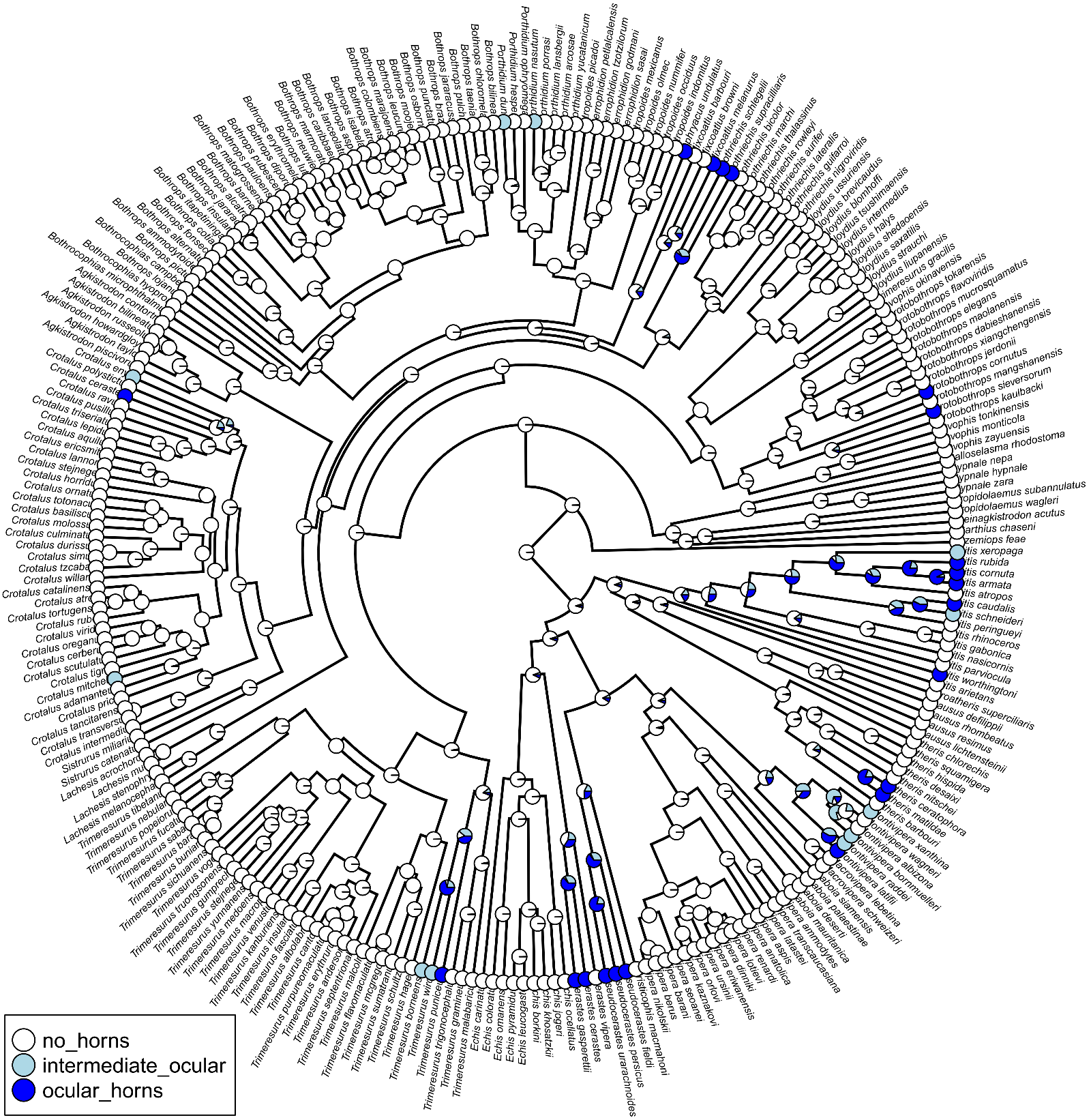

**Figure S5.** Ancestral state reconstruction of ocular horns within the Viperidae. Pies depict the posterior probability of character states at each node obtained from 1000 stochastic character maps.

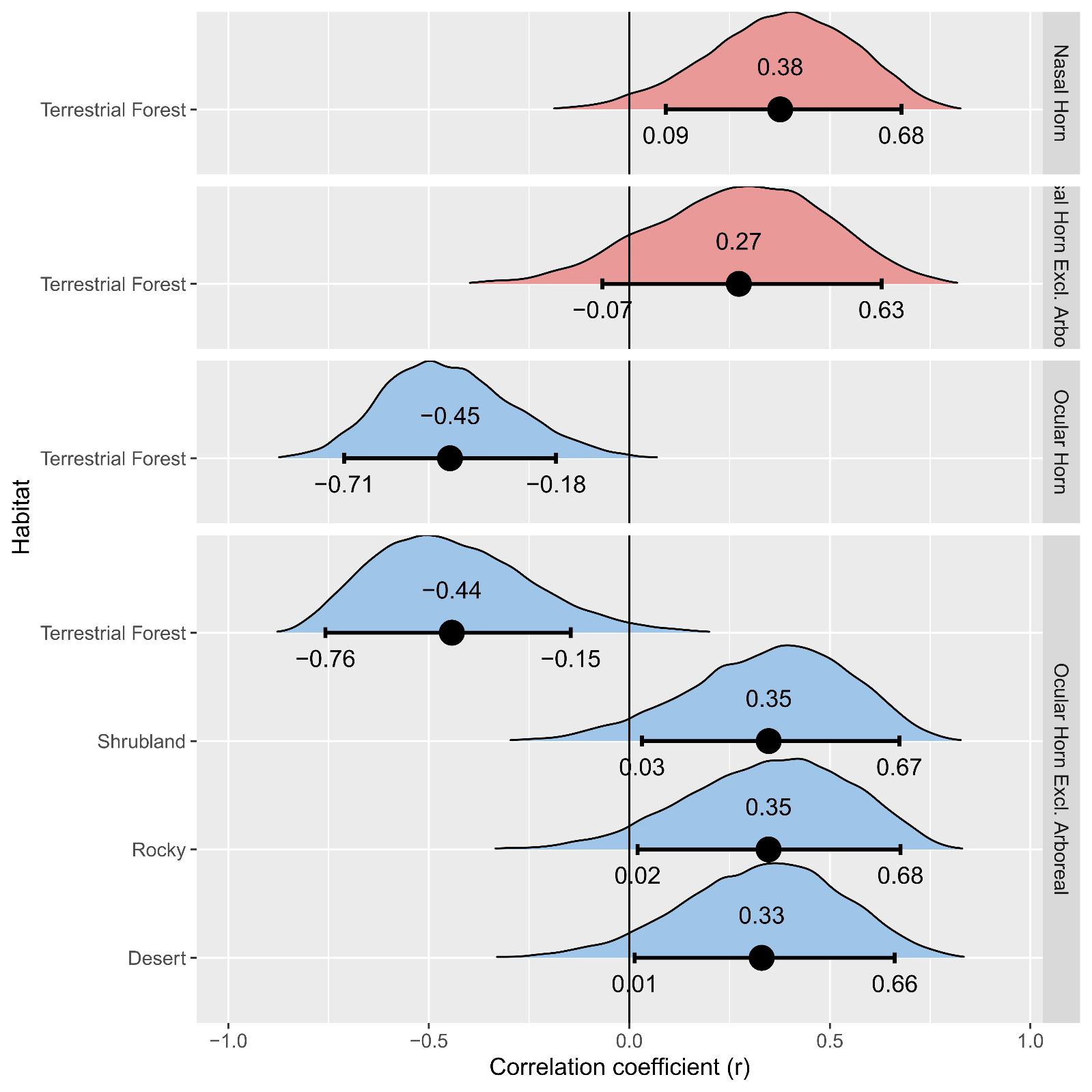

**Figure S6.** Posterior estimate of the correlation coefficient (r) between the presence of horns and habitat. Showing results for nasal (red) and ocular (blue) horns including all taxa and when arboreal taxa were excluded. Dot indicates the mean of the posterior distribution. Error bars indicate the 90% highest posterior density (HPD) to assess the probability of direction (PD). Direction of the correlation is regarded as significant where the 90% HPD excludes zero, i.e., the PD is >95% positive or negative.

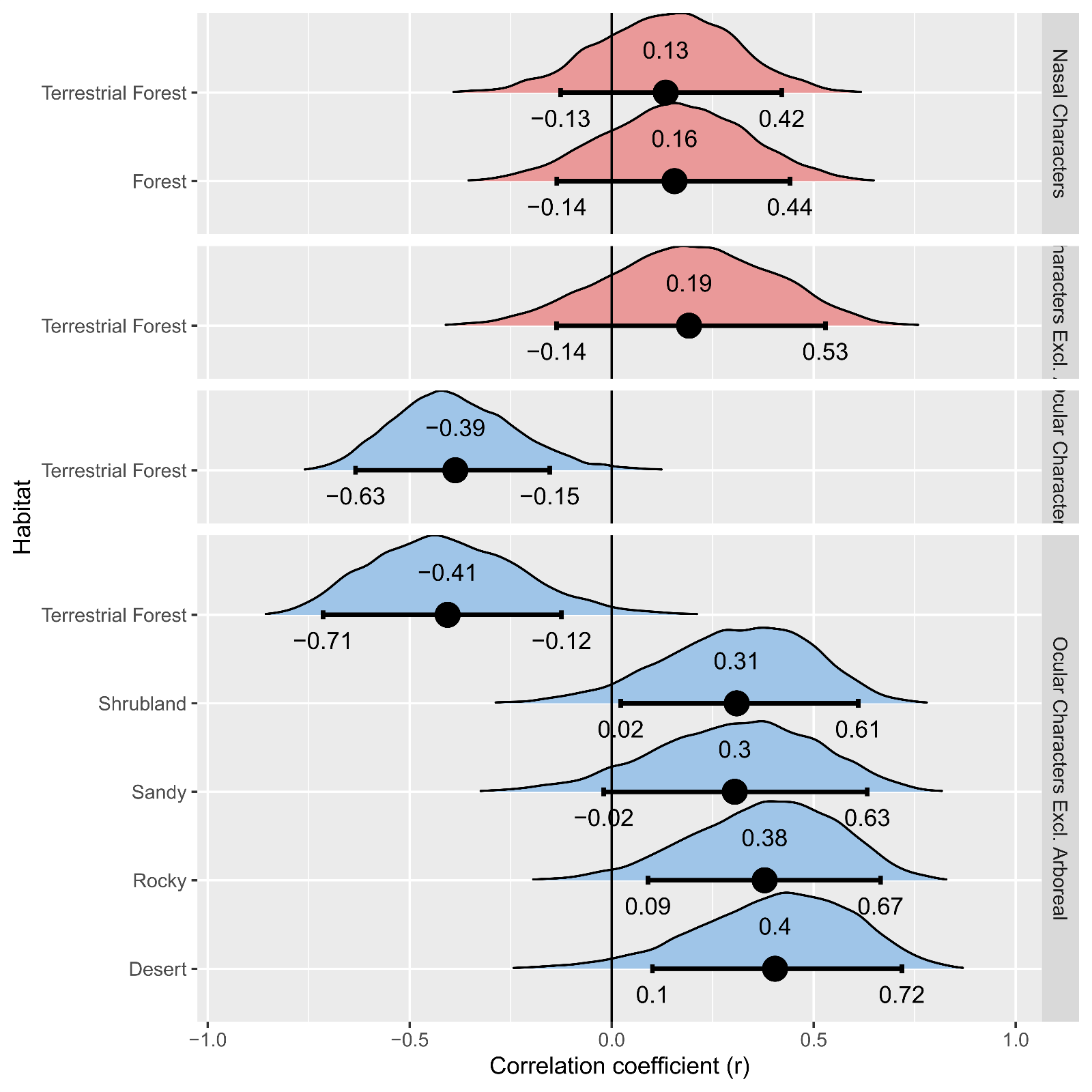

**Figure S7.** Posterior estimate of the correlation coefficient (r) between the presence of characters and habitat, where characters refer to the combination of intermediate and horned states. Showing results for nasal (red) and ocular (blue) characters including all taxa and when arboreal taxa were excluded. Dot indicates the mean of the posterior distribution. Error bars indicate the 90% highest posterior density (HPD) to assess the probability of direction (PD). Direction of the correlation is regarded as significant where the 90% HPD excludes zero, i.e., the PD is >95% positive or negative.

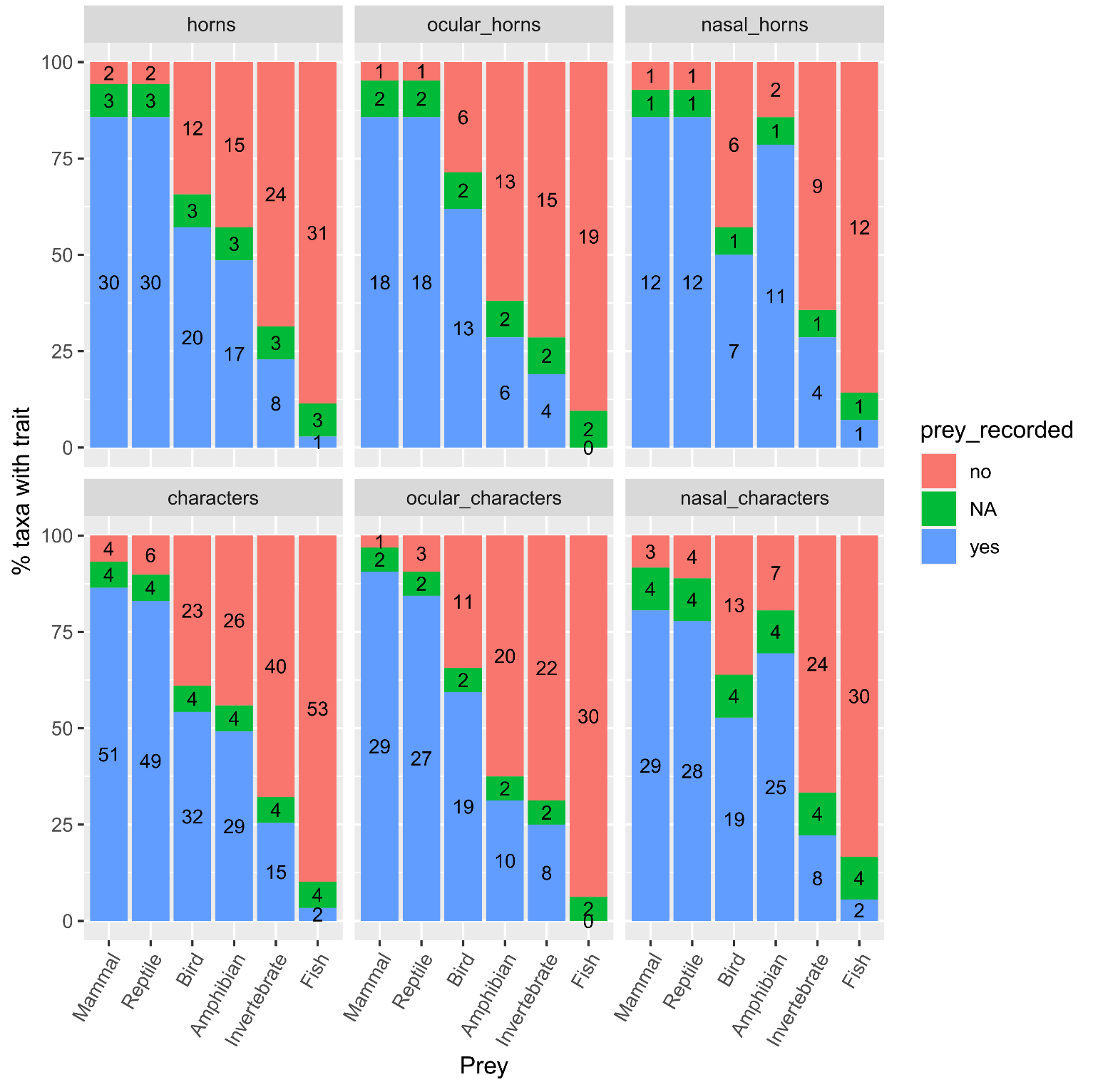

**Figure S8.** Relative proportion of Viperidae taxa with cephalic traits recorded feeding on each prey class. Values in the columns reflect the absolute number of taxa. NA depicts taxa for which there is no diet information.
